# Somatic evolution of a cross-reactive germline antibody that expands its breadth to neutralize new SARS-CoV-2 variants

**DOI:** 10.64898/2026.03.20.713312

**Authors:** Huibin Lv, Ziqi Feng, Qi Wen Teo, Chunke Chen, Akshita B. Gopal, Danbi Choi, Timothy J. C. Tan, Yun Sang Tang, Lewis Siu, Armita Nourmohammad, Roberto Bruzzone, Ian A. Wilson, Meng Yuan, Nicholas C. Wu, Chris K. P. Mok

## Abstract

Rapid antigenic drift of the SARS-CoV-2 receptor-binding domain (RBD) underlies immune escape and continues to challenge the durability of antibody-mediated protection. Among the major classes of RBD-directed antibodies, germline-encoded *IGHV3-53* responses are highly potent against early SARS-CoV-2 variants but are generally compromised by Omicron-associated mutations. Here, we identify an intrinsically cross-reactive *IGHV3-53* germline antibody that recognizes multiple pre-Omicron variants, including SARS-CoV-2 wild-type, Alpha, and Delta. Notably, we demonstrate that targeted somatic evolution can further expand this breadth to overcome the immune escape of different Omicron variants. Guided by integrated structural and sequence analyses, we introduce four somatic mutations (G26E, T28I, S53P, and Y58F) into the germline antibody, resulting in markedly enhanced binding and neutralization of Omicron BA.1, BA.2, and BA.4/5. High-resolution crystal structures reveal that these mutations re-establish critical interactions disrupted by substitutions on Omicron RBD and optimize affinity at a remodeled epitope interface. Collectively, our findings delineate a structural and mechanistic pathway through which an inherently cross-reactive germline antibody lineage can be adaptively refined to counter highly divergent SARS-CoV-2 variants. This work highlights the underappreciated breadth encoded within the naïve B-cell repertoire and provides a conceptual framework for engineering and eliciting antibody responses resilient to future antigenic drift.

## INTRODUCTION

The continual emergence of SARS-CoV-2 variants of concern (VOCs), including Alpha, Beta, Gamma, Delta, and multiple antigenically divergent Omicron sublineages (e.g., BA.1, BA.5, XBB.1.5, JN.1, and KP.3),^1^ has posed a persistent challenge to global public health and economic stability. Mutations accumulating in the viral spike protein, particularly within the receptor-binding domain (RBD), enable evasion of pre-existing neutralizing antibodies elicited by vaccination or prior infection, thereby enhancing viral transmissibility and undermining population immunity.^2,3^ This antigenic evolution has also substantially reduced the effectiveness of most therapeutic monoclonal antibodies, a problem that became especially pronounced following the global spread of Omicron variants.^1,4,5^ Consequently, defining how broadly neutralizing antibodies (bnAbs) achieve and maintain breadth against rapidly evolving variants has become a central objective in combating SARS-CoV-2 evolution.

Upon antigen encounter, B cells undergo affinity maturation through the accumulation of somatic hypermutations (SHMs) that enhance antibody affinity and specificity.^6^ Studies showed that this affinity maturation process can last for up to 1 year after infection of SARS-CoV-2 or immunization with COVID-19 vaccine.^7-13^ Interestingly, it has been suggested that affinity maturation of some antibodies that are specifically against the ancestral SARS-CoV-2 strain can increase the neutralization breadth against the new VOCs.^8,11,14^ For instance, while the spike protein of wild-type SARS-CoV-2 with Q493R and E484G mutations was resistant to viral neutralization by monoclonal antibody (mAb) C144, another mAb from the same clonotype with different SHMs, C501, could potently neutralize these mutant strains.^8^ Moreover, receiving three doses of monovalent ancestral wild-type SARS-CoV-2 mRNA vaccine could induce antibodies against the Omicron BA.1 and BA.2, which these newly appeared Omicron variants are antigenically distinct from the wild-type strain.^15-17^ However, the mechanism used during affinity maturation that leads to increased cross-reactivity of B cells to overcome viral escape remains largely elusive.

*IGHV3-53/66* antibodies constitute a major and recurrent class of nAbs elicited by SARS-CoV-2 infection or vaccination with pre-Omicron strains.^18^ Although the majority of pre-existing antibodies to wild-type are evaded, the persistence of rare *IGHV3-53/66* antibodies with substantial activity against Omicron variants raises a critical question as to how resistance to antigenic drift can be achieved. Here, we characterize a germline-encoded *IGHV3-53* antibody HB148, devoid of SHM, isolated from an unvaccinated individual infected with SARS-CoV-2 during the first wave of the COVID-19 outbreak. This antibody neutralizes multiple pre-Omicron variants but lacks activity against Omicron sublineages due to mutations on the RBD. Four SHMs, which were identified by sequence analyses, were engineered into HB148, and neutralizing activity to different Omicron strains was examined. Furthermore, we determined high-resolution crystal structures to understand the mechanisms by which a germline precursor can acquire mutations to counteract the evolution of newly emerging variants.

## Results

### A neutralizing germline antibody against SARS-CoV-2 RBD

In our previous study,^19^ we employed biotinylated SARS-CoV-2 RBD to enrich wild-type RBD-specific B cells from PBMC samples collected from 19 individuals with confirmed SARS-CoV-2 infection, followed by single B cell sequencing. These samples were obtained within 40 days of symptom onset, capturing the early phase of the humoral immune response. From the screening, we identified a germline-encoded antibody HB148, which binds to the full-length spike and RBD proteins of the ancestral SARS-CoV-2 **(Figure S1A)**. HB148 exhibited potent binding to SARS-CoV-2 RBD, with an EC_50_ of 150 ng/mL **(Figure S1B)**. HB148 is encoded by *IGHV3-53/IGKV3-20* and no SHM was detected in either its heavy or light chain **(Figure S1C)**, indicating that HB148 is a germline antibody that may function as an early defense against SARS-CoV-2 infection.

Next, we used the surrogate virus neutralization test (sVNT) to further evaluate the cross-neutralizing potency of HB148. Of note, the surrogate virus neutralization test (sVNT) has been widely utilized to assess the neutralizing capacity of SARS-CoV-2 in serum and plasma samples from COVID-19 convalescents^20^, since it positively correlates with those from the live virus neutralization assay.^21^ HB148 was tested against the SARS-CoV-2 wild-type strain as well as five VOCs: Alpha (B.1.1.7), Beta (B.1.351), Gamma (P.1), Delta (B.1.617.2), and Omicron BA.1 (B.1.1.529) **(Figure 1A-F)**. Compared to the wild-type strain (NT_50_ = 1.88 μg/mL), HB148 demonstrated comparable neutralization potency against Alpha (NT_50_ = 1.00 μg/mL) and Delta (NT_50_ = 1.39 μg/mL), but reduced potency against Beta (NT_50_ = 75.99 μg/mL) and Gamma (NT_50_ = 10.55 μg/mL) variants **(Figure 1A-E)**. However, HB148 failed to neutralize the Omicron BA.1, even at the highest tested concentration of 100 μg/mL **(Figure 1F)**. To further validate the neutralization results, we used a SARS-CoV-2 pseudovirus system. Generally consistent with the sVNT findings, HB148 exhibited strong neutralizing activity against the pseudoviruses of wild-type (NT_50_ = 7.39 μg/mL), Alpha (NT_50_ = 0.50 μg/mL), and Delta (NT_50_ = 0.06 μg/mL), while showing weak activity against Beta strains (NT_50_ = 27.21 μg/mL). Notably, no neutralizing activity was observed against the Gamma or Omicron pseudoviruses, even at 100 μg/mL **(Figure S2A-S2F)**. These findings are consistent with previous studies,^4,22,23^ showing that mutations acquired by the Omicron variant BA.1 enable escape from most antibodies elicited by infection or vaccination with the SARS-CoV-2 wild-type strain.

**Figure 1.**
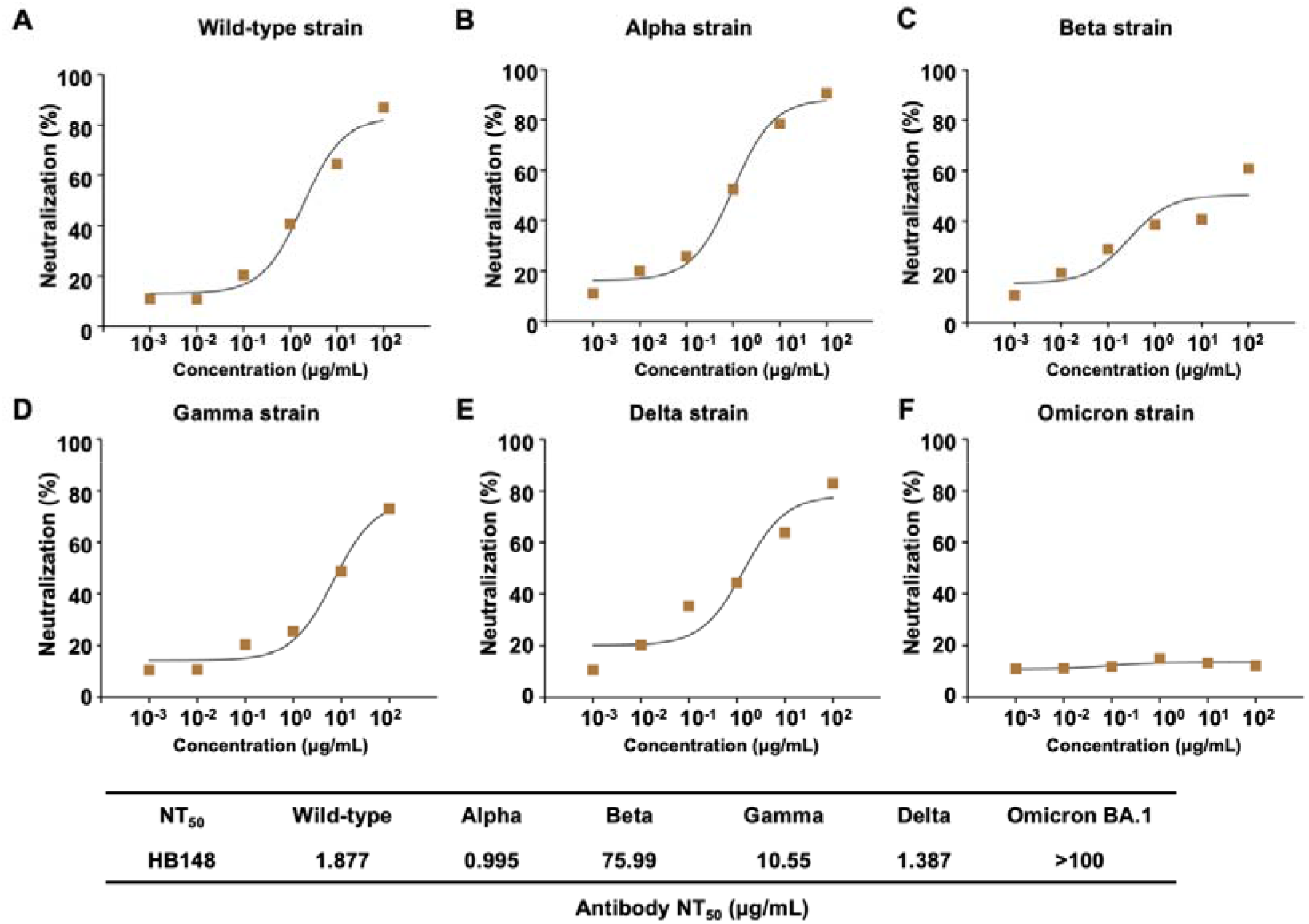
Antibody neutralization against SARS-CoV-2 wild-type and variants by sVNT. Neutralization titer 50 (NT_50_) of antibody HB148 against **(A)** wild-type strain, **(B)** Alpha strain, **(C)** Beta strain, **(D)** Gamma strain, **(E)** Delta strain and **(F)** Omicron BA.1 strain were measured by surrogate virus neutralization test (sVNT). Their NT_50_ values are indicated.

### Structural analysis of germline HB148 with wild-type RBD

To define the antibody binding site, RBD epitope, and elucidate the structural basis of HB148-mediated cross-neutralization, we determined a crystal structure of HB148 in complex with the SARS-CoV-2 wild-type RBD at 3.04 Å resolution **(Table S1)**. Similar to other *IGHV3-53* antibodies,^24-26^ HB148 targets the class 1 epitope on the RBD **(Figure 2A)**. The interaction is dominated by the heavy chain, which contributes 69% of the buried surface area (BSA), while the light chain accounts for the remaining 31%. HB148 engages several conserved residues shared among the wild-type and variant RBDs, including Y421, Y453, and A475 **(Figure 2B and Figure S3)**.

**Figure 2.**
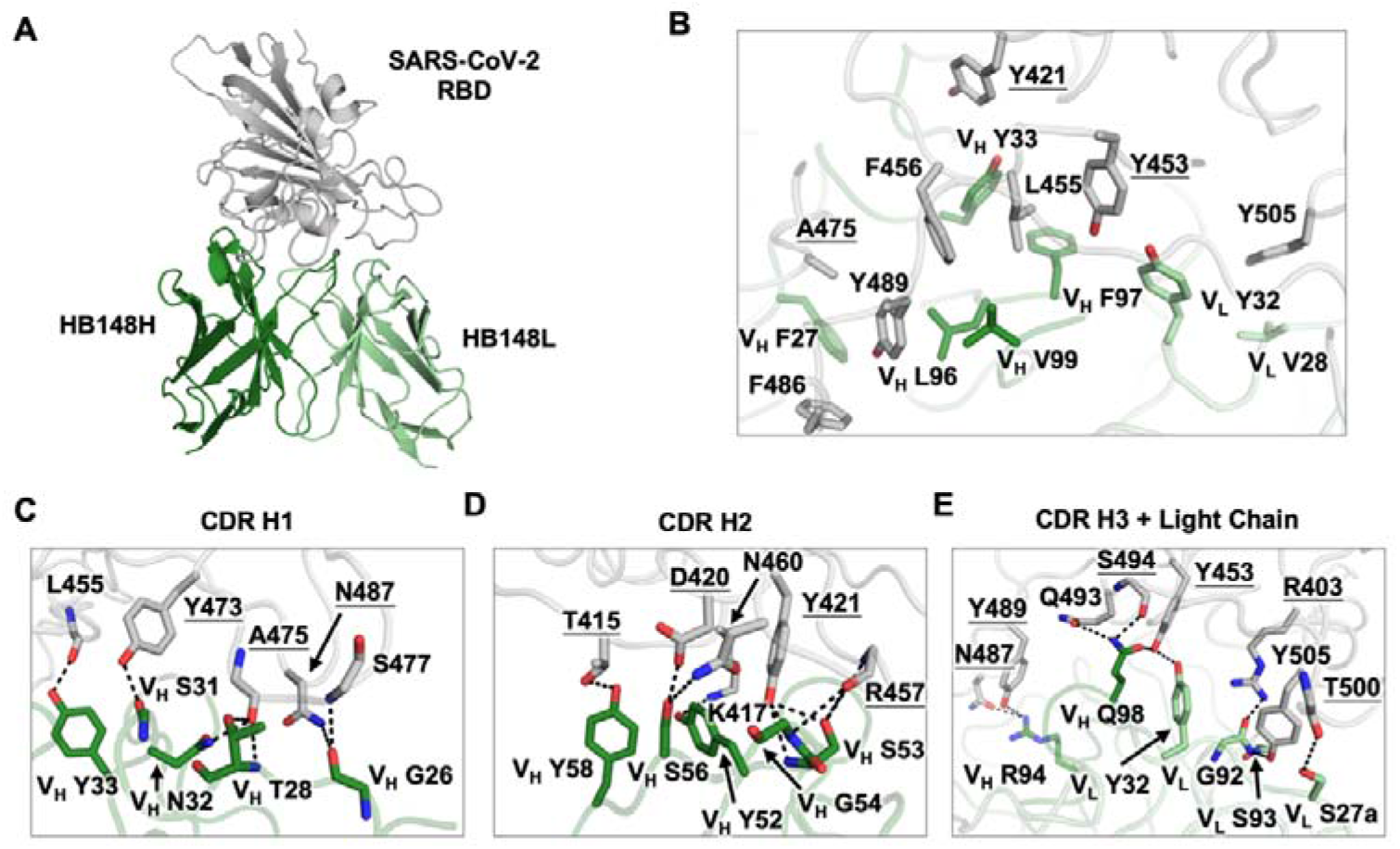
Structural analysis of HB148 with wild-type RBD. **(A)** X-ray crystal structure of HB148 antibody in complex with SARS-CoV-2 wild-type RBD. **(B)** Hydrophobic interactions between HB148 and SARS-CoV-2 wild-type RBD. Kabat numbering is applied to the antibody. Conserved epitope residues across VOCs are underlined. Detailed interactions (H-bonds and salt bridges) of SARS-CoV-2 RBD with HB148 are shown in **(C)** CDR H1, **(D)** CDR H2, and **(E)** CDR H3+Light Chain respectively.

Within the paratope, HB148 CDR H1 forms a compact hydrogen-bonding network with the wild-type RBD **(Figure 2C)**, where V_H_ N32, T28, and G26 hydrogen bond with conserved residues Y473, A475, and N487, respectively. CDR H2 contributes additional contacts through V_H_ Y58 and S56, which interact with T415 and D420 on the RBD **(Figure 2D)**. CDR H3, of moderate length (14 residues), contributes substantially to antigen engagement by adopting an extended conformation that cooperates with the light chain. Specifically, V_H_ Q98 and R94 within CDR H3 act together with light-chain residues V_L_ S93, G92, and S27a to anchor HB148 to key RBD hotspots, including Y489, R403, and T500, through a dense hydrogen-bonding network and stabilizing hydrophobic stacking interactions.

To further understand how viral variants escape HB148 recognition, we aligned RBD sequences from representative SARS-CoV-2 variants **(Figure S3)**. The RBD residues contacted by HB148, including T415, K417, R457, A475, S477, and N487, are conserved among wild-type, Alpha, and Delta variants, consistent with the comparable sVNT neutralizing activities observed against these strains **(Figure 1A-1B and 1E)**. In the Beta and Gamma variants, the K417N/T substitutions likely disrupt polar interactions with V_H_ S53 and G54, thereby reducing neutralizing activity **(Figure 1C-1D)**. In Omicron BA.1, multiple substitutions, K417N, S477N, Q493R, and Y505H, collectively remodel the HB148-binding surface on the RBD and completely abrogate binding **(Figure 1F)**. These alterations are consistent with prior structural and functional studies^27-29^ showing that these mutations mediate broad escape from *IGHV3-53*–derived and other class 1 neutralizing antibodies.

### Antibody signature analysis of the *IGHV3-53* antibody

Several *IGHV3-53*–derived antibodies have been reported to retain binding and neutralizing activity against the Omicron BA.1 and later emerged variants,^29,30^ suggesting that members of this germline family can acquire cross-reactivity through SHMs. To investigate the molecular features underlying Omicron BA.1 recognition, we analyzed *IGHV3-53* antibodies from the Coronavirus Antibody Database (CoV-AbDab)^31^ with available sequence and neutralizing data. Compared with non–BA.1-neutralizing antibodies, BA.1-neutralizing antibodies exhibited an enrichment of four SHMs: V_H_ G26E and T28I in CDR H1, as well as V_H_ S53P and Y58F in CDR H2 **(Figure 3A and Figure S4)**. Different combinations of these mutations were introduced into HB148, and binding affinities to wild-type and BA.1 RBDs were measured using biolayer interferometry (BLI). Notably, incorporation of all four mutations (HB148-M4) resulted in an approximately 100-fold decrease in K_D_ for both the wild-type and BA.1 RBDs, indicating a marked enhancement in binding affinity **(Figure S5)**. Moreover, single mutation for V_H_ G26E, S53P and Y58F individually decreased the K_D_ for BA.1 RBD binding by approximately 8- to 10-fold **(Figure S5)**, suggesting that these substitutions act additively to optimize paratope–epitope complementarity and promote cross-reactivity toward Omicron BA.1.

**Figure 3.**
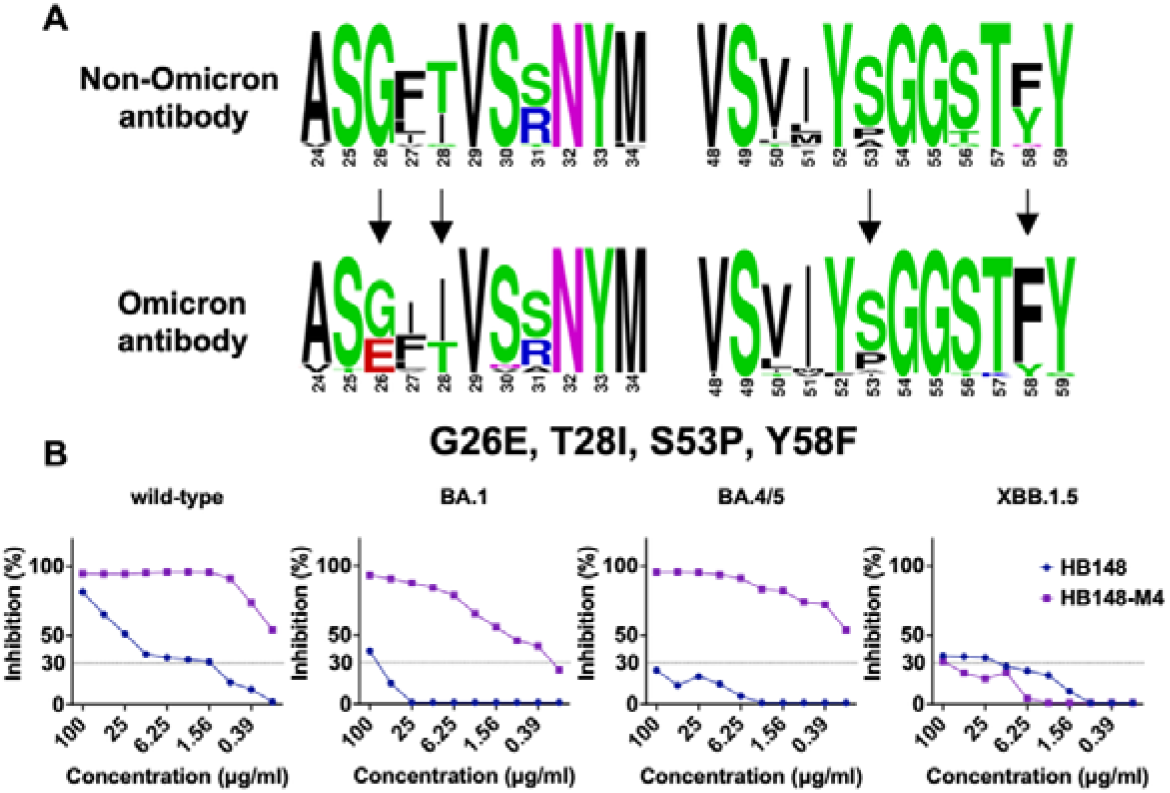
Four somatic hypermutations restore binding of HB148 to Omicron strains. **(A)** Sequence alignment between non-Omicron neutralizing antibodies and Omicron neutralizing antibodies. Sequences are downloaded from the Coronavirus Antibody Database (CoV-AbDab). Sequence logo has been plotted by WebLogo (https://weblogo.berkeley.edu/logo.cgi). **(B)** Neutralization titer 50 (NT_50_) of antibody HB148 and HB148-M4 against SARS-CoV-2 wild-type strain, Omicron BA.1 strain, BA.4/5 strain and XBB.1.5 strain were measured by surrogate virus neutralization test (sVNT).

We next assessed the neutralizing activity of HB148 variant against SARS-CoV-2 wild-type, BA.1, BA.4/5, and XBB.1.5 using sVNT assays. Compared to HB148, HB148-M4 neutralized SARS-CoV-2 wild-type, BA.1, and BA.4/5 pseudoviruses, with NT_50_ values ranging from 0.19 to 1.56 μg/mL **(Figure 3B)**. In contrast, neither HB148 nor HB148-M4 neutralized XBB.1.5. These results demonstrate that the four mutations substantially enhance HB148’s capacity to overcome immune escape by Omicron BA.1 and BA.4/5, whereas additional substitutions in XBB.1.5 further remodel the RBD surface and abrogate antibody recognition, implying that additional paratope adaptations would be required to restore binding.

### Structural analysis of HB148-M4 with wild-type RBD and BA.1 RBD

To determine how HB148-M4 gains binding affinity to overcome the immune escape by Omicron strains, we determined crystal structures of HB148-M4 in complex with the SARS-CoV-2 wild-type RBD and Omicron BA.1 RBD at 2.60 Å and 2.73 Å resolutions, respectively **(Figure 4A-4D and Table S1)**. The heavy chain of HB148-M4 contributes 58% and 57% of the BSA to the wild-type and BA.1 RBDs, respectively, whereas the light chain accounts for the remaining 42% and 43%.

**Figure 4.**
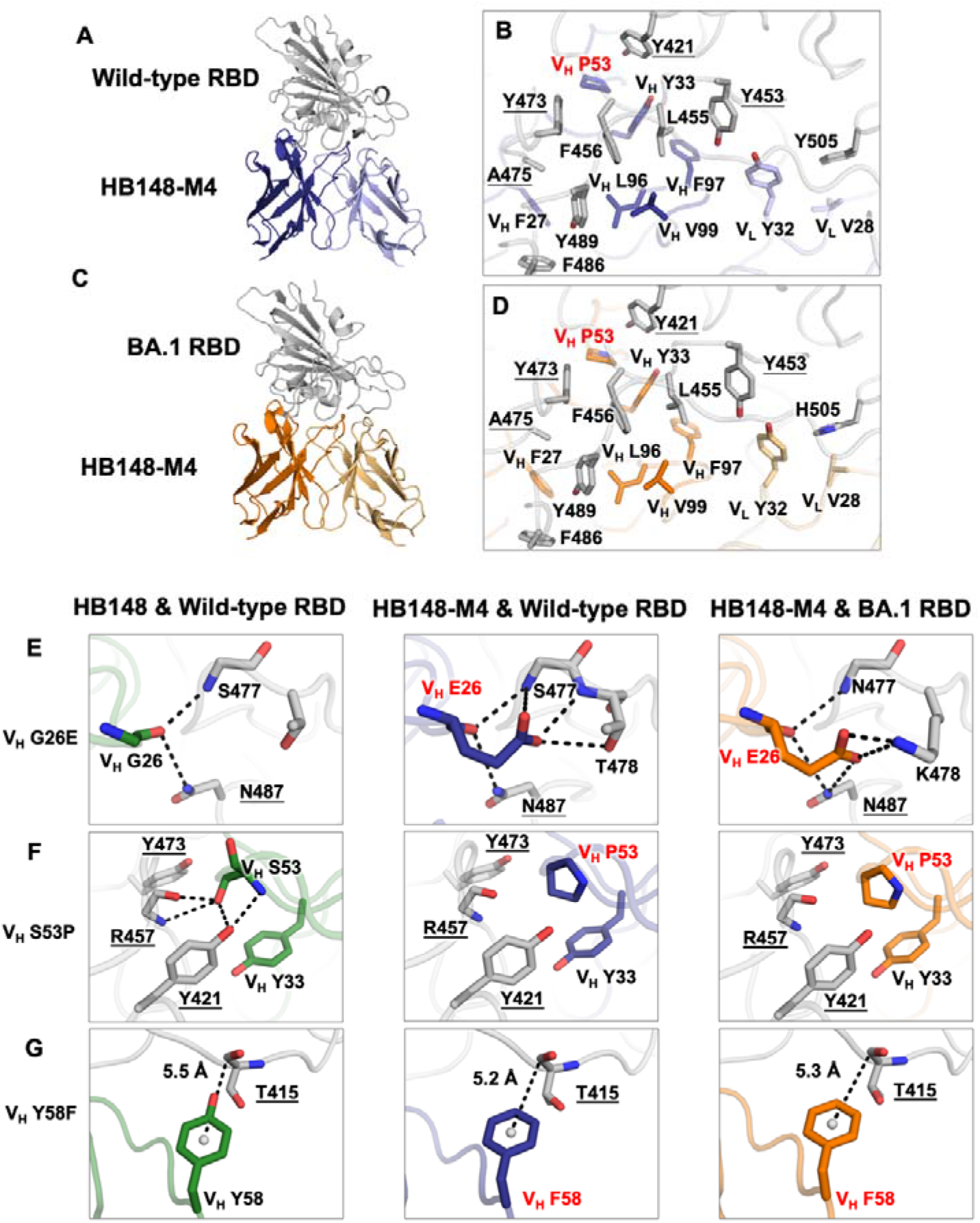
Structure analysis of HB148-M4 with SARS-CoV-2 wild-type RBD and BA.1 RBD. **(A, C)** X-ray crystal structure of HB148-M4 antibody in complex with RBD from **(A)** SARS-CoV-2 wild-type and **(C)** BA.1 RBD. **(B, D)** Hydrophobic interactions between HB148-M4 and **(B)** SARS-CoV-2 wild-type RBD and **(D)** BA.1 RBD. Kabat numbering is applied to the antibody. Conserved epitope residues across VOCs are underlined. Four mutations (V_H_ G26E, T28I, S53P, and Y58F) are labeled in red. Detailed interactions (H-bonds and salt bridges) of RBD with HB148-M4 are shown with **(E)** V_H_ G26E, **(F)** V_H_ S53P, and **(G)** V_H_ Y58F, respectively.

Specifically, the V_H_ T28I mutation exhibited similar interactions with N475 of both wild-type and BA.1 RBDs as V_H_ T28 did with the wild-type RBD **(Figure S6)**, consistent with the observation that this single substitution did not enhance binding affinity for either RBD **(Figure S6)**. In contrast, substitution of V_H_ G26 with E26 in CDRH1 introduced additional hydrogen bonds and enabled formation of salt bridges with K478 in the BA.1 RBD **(Figure 4E)**. Additionally, replacement of V_H_ S53 with P53 in CDRH2 disrupted hydrogen bonds with Y421 and R457 but established hydrophobic interactions with V_H_ Y33, Y421, and Y473 **(Figure 4F)**. Furthermore, substitution of V_H_ Y58 with F58 in CDRH2 positioned the residue closer to the backbone amide of RBD T415, strengthening T-shaped π–π stacking interactions **(Figure 4G)**, which is consistent with our previous finding.^32^ Collectively, these structural features suggest that the V_H_ G26E, S53P, and Y58F mutations contribute to enhanced binding affinity toward both wild-type and BA.1 RBDs.

The Omicron BA.1 RBD primarily uses K417N, S477N, Q493R, and Y505H substitutions to evade HB148 binding **(Figure 2C-2E)**. Specifically, K417N likely disrupts interactions with V_H_ Y52 in CDRH2, Q493R alters contacts with V_H_ Q98 in CDRH3, and Y505H interferes with interactions involving V_L_ G92 in CDRL3. The four mutations introduced into HB148 alone could not fully restore binding to these altered sites. However, the V_H_ G26E substitution extends interactions between V_H_ E26 and RBD residues 477 and 478, facilitating partial restoration of binding to the BA.1 RBD. Additionally, the V_H_ S53P and Y58F double mutations increased binding affinity toward both wild-type and BA.1 RBDs **(Figure S5)**. Together, these results suggest that the combined effects of the four substitutions in HB148-M4 may additively enhance binding and neutralization potency against the Omicron BA.1 variant.

### Extend the four mutations to other *IGHV3-53* antibodies

To determine whether the four SHMs could enhance the cross-reactivity of other *IGHV3-53* antibodies, we introduced them into C1A-F10,^33^ P5A-3C8,^26^ BG4-25,^34^ and LY-CoV488.^35^ All four antibodies bound to the wild-type RBD but failed to or weakly recognize the BA.1 RBD. Incorporation of the four mutations restored BA.1 binding for P5A-3C8, BG4-25, and LY-CoV488, but not C1A-F10, although C1A-F10-M4 showed increased affinity for wild-type RBD **(Figure 5A)**. Consistent with the binding data, the M4 version also exhibited extended neutralizing activity against the Omicron BA.1 and BA.4/5 strains **(Figure 5B)**. However, all mutants showed minimal neutralization against the Omicron XBB.1.5 strain, even at 100 μg/mL **(Figure 5B)**. These findings suggest that the M4 mutations can broaden the binding and neutralization breadth of *IGHV3-53* antibodies against early Omicron variants.

**Figure 5.**
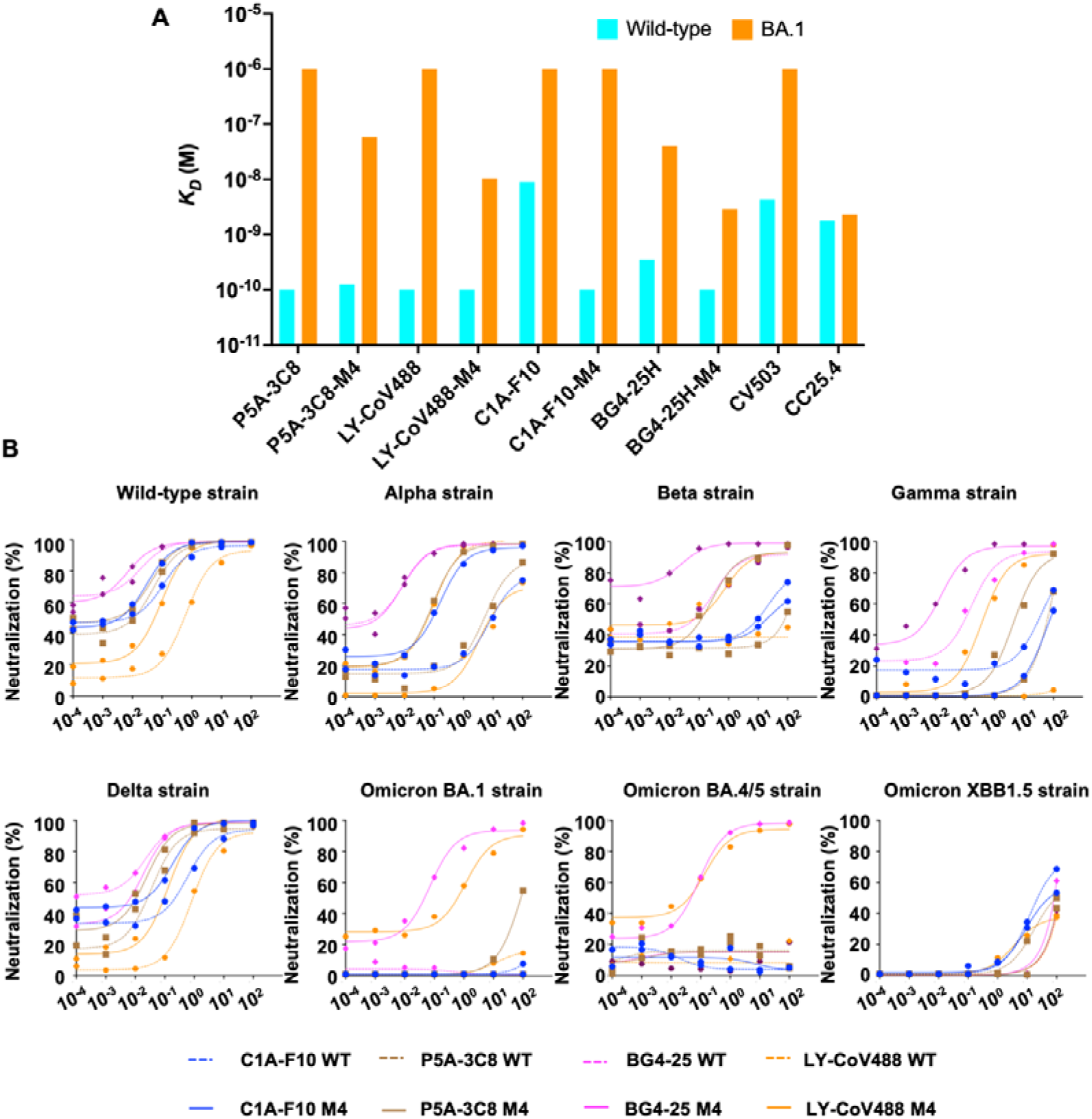
Four somatic hypermutations restore binding for other *IGHV3-53* antibodies. **(A)** Binding kinetics of *IGHV3-53* antibodies against wild-type RBD and BA.1 RBD were measured by BLI. Y-axis represents the response. CV503,^58^ a wild-type RBD–binding antibody, and CC25.4,^59^ which binds both wild-type and BA.1 RBDs, were used as controls. **(B)** NT_50_ of antibody HB148 and HB148-M4 against SARS-CoV2 wild-type strain, Omicron BA.1 strain, BA.4/5 strain, and XBB.1.5 strain was measured by sVNT.

## Discussion

In this study, we identified an *IGHV3-53* germline-encoded antibody, HB148, that neutralizes multiple SARS-CoV-2 variants in the absence of SHM. During the early stage of the pandemic, most isolated SARS-CoV-2-specific mAbs had few SHMs.^36-42^ Other germline antibodies have shown similar activity, underscoring the latent capacity of the naïve human B cell repertoire to recognize novel pathogens.^43^ However, the rapid antigenic evolution of SARS-CoV-2 enables escape from this baseline immunity, requiring SHM to broaden antibody breadth. Using sequence and structural analyses, we delineate a single cycle of viral escape and antibody adaptation, illustrating how mutations in the virus and host antibody reciprocally shape each other in this ongoing molecular arms race.

Notably, the four mutations did not confer breadth to all *IGHV3-53* antibodies. For example, in C1A-F10, the M4 substitutions increased affinity for the wild-type RBD but failed to increase BA.1 binding, indicating that enhanced affinity alone is insufficient for some antibodies to increase recognition of BA.1. BLI and structural analyses of HB148 with wild-type and BA.1 RBD suggest two mechanisms for restoring BA.1 binding: increasing overall affinity and establishing new contacts with BA.1-specific residues. Thus, antibodies that cannot regain BA.1 binding may be limited by features of their paratope, such as light-chain-dependent contacts or may require additional SHMs to accommodate the altered BA.1 epitope.

Furthermore, the M4-substituted antibodies showed minimal neutralization of the XBB.1.5 strain. XBB.1.5 exhibits near-complete escape from neutralizing antibody responses elicited by three doses of mRNA vaccine or by BA.4/5 infection.^44^ Compared with BA.4/5, XBB.1.5 accumulates additional spike mutations, including R346T, L368I, V445P, and S486P within the RBD. Notably, F486 is a key antigenic hotspot for class 1 neutralizing antibodies. P5A-3C8, BG4-25, and LY-CoV488 engage F486 through a hydrophobic cage formed by multiple paratope residues. This interaction is disrupted by the F486S/P substitutions in BA.4/5 and XBB.1.5. Interestingly, although BA.4/5 carries F486V, a non-aromatic residue, P5A-3C8-M4 and BG4-25-M4 still neutralized BA.4/5, suggesting that loss of the aromatic ring alone does not fully explain escape. These observations indicate that additional XBB.1.5-specific RBD mutations act in combination to evade recognition by the M4-modified antibodies.

*IGHV3-53/66* antibodies represent one of the most frequently elicited classes of neutralizing antibodies following SARS-CoV-2 infection or vaccination with pre-Omicron strains, but they are largely evaded by Beta, Gamma, and Omicron variants. Several studies have investigated the molecular determinants of *IGHV3-53/66* antibody clones with the capacity to give rise to rare evasion-resistant antibodies, such as BD55-1205 and V30V4.^30,45-47^ In addition to the heavy-chain V gene usage, light-chain pairing and CDR H3 length also appear to play important roles in shaping the potential of these clones to further evolve toward bnAbs. Consistent with these observations, our study demonstrates that hallmark somatic mutations play a critical role in enhancing both the neutralization potency and breadth of *IGHV3-53/66* antibodies.

Overall, our study highlights the ongoing molecular tug of war between SARS-CoV-2 evolution and the host antibody response. As the virus accumulates mutations to evade recognition, immunity is continually reshaped through vaccination, infection, and antibody maturation. By dissecting a full cycle of viral escape and antibody adaptation, we provide a proof-of-concept framework for engineering cross-reactive antibodies, offering valuable insights for future therapeutic antibody and vaccine design.

## Materials and methods

### Cells

The Expi293F cells were cultured in Expi293™ Expression Medium and passaged every 3 to 4 days. The HEK293T and A549-ACE2 stable cell lines were cultured in Dulbecco’s modified Eagle’s medium (DMEM high glucose; Gibco) supplemented with 10% heat-inactivated fetal bovine serum (FBS; Gibco), 1% penicillin-streptomycin (Gibco) and 1% Gluta-Max (Gibco). Of note, A549-ACE2 stable cell lines were selected by using Puromycin (5 μg/mL). Sf9 cells (*Spodoptera frugiperda* ovarian cells, female, ATCC) were maintained in Sf-900 II SFM medium (Thermo Fisher Scientific).

### Expression and purification of human mAbs

B cells were selected for mAb generation based on RBD antigen probes, as previously described.^11,42^ Antibody heavy and light chain genes obtained by 10X Genomics V(D)J sequencing analysis was synthesized by IDT (Integrated DNA Technologies). The synthesized fragments for heavy and light chains with 5’ and 3’ Gibson overhangs were then cloned into human IgG1 and human kappa or light chain expression vectors by Gibson assembly as previously described.^48^ Plasmids encoding heavy and light chains were transfected into Expi293F cells in a 2:1 mass ratio using an ExpiFectamine 293 transfection kit (Gibco). Six days post-transfection, the supernatant was harvested and purified using CaptureSelect CH1-XL beads (Thermo Scientific). The purified antibodies were boiled and subjected to SDS-PAGE gel for Coomassie Blue R-250 staining.

### Expression and purification of Fab proteins

The heavy and light chains of the Fab were cloned into the phCMV3 vector. The plasmids were transiently co-transfected into ExpiCHO cells at a ratio of 2:1 (heavy chain to light chain) using ExpiFectamine CHO Reagent (Thermo Fisher Scientific) according to the manufacturer’s instructions. The supernatant was collected at 7 days post-transfection. The Fab was purified with a CaptureSelect CH1-XL Pre-packed Column (Thermo Fisher Scientific), followed by SEC and buffer exchanged 20□mM Tris-HCl, 150 mM NaCl, pH 7.4.

### Enzyme-linked immunosorbent assay

Nunc MaxiSorp ELSIA plates (Thermo Fisher Scientific) were coated with 100 μL of recombinant proteins at 1 □μg/mL in a 1× PBS solution overnight at 4°C. The plates were washed 3 times the next day with 1× PBS supplemented with 0.05% Tween 20 and blocked with 200 □μL of 1× PBS containing 5% non-fat milk powder for 2 □h at room temperature. mAbs were serially diluted 1:10 starting at 100 □μg/mL and incubated for 2 □h at 37°C. The plates were then washed 3 times and incubated with horseradish peroxidase (HRP)-conjugated goat anti-human IgG antibody (GE Healthcare) diluted 1:5,000 for 1 □h at 37°C. The ELISA plates were then washed five times with PBS containing 0.05% Tween 20. Subsequently, 100 μL of TMB solution (Sigma) was added into each well. After 15 min of incubation, the reaction was stopped by adding 50 μL of 2 □M H_2_SO_4_ solution and analyzed on a Sunrise (Tecan, Männedorf, Switzerland) absorbance microplate reader at 450 nm wavelength.

### Surrogate Virus Neutralization Test (sVNT)

SARS-CoV-2 wild-type and the VOCs sVNT (cPass) was performed according to the manufacturer’s instructions.^21^ In brief, HRP-RBD conjugate was pre-incubated with serially 10-fold diluted antibodies (starting at 100□μg/mL) for 30 min followed by addition to the ACE2-coated ELISA plate for 15 min. After four washes, the colorimetric signal was developed by TMB substrate and 2□M H_2_SO_4_ stop solution. Absorbance reading at 450 nm were acquired using Sunrise (Tecan, Männedorf, Switzerland) absorbance microplate reader.

### Production of SARS-CoV-2 Spike pseudoviruses

The pNL Luc E-R-vector encoding the luciferase reporter gene was previously used in HIV/MERS spike pseudo-particles construction.^49^ To generate the SARS-CoV-2 Spike and viral mutants pseudotyped HIV-1 single-round luciferase virus, HEK293T cells were co-transfected with pNL4-3. Luc R-E™ and recombinant SARS-CoV-2 Spike plasmids (pcDNA-SARS-CoV-2 spike) using the PEI transfection reagent (Polysciences) according to the manufacturer’s instructions. At 72 h post transfection, collect virus by harvesting the supernatant from each well and filtering it through a 0.45 μm SFCA low protein-binding filter. Virus can be stored at 4 °C for immediate use or frozen at ™80 °C.

### Pseudotyped virus neutralization assay

Serially diluted monoclonal antibodies were incubated with SARS-CoV-2 pseudotyped virus for 1 h at 37 □°C. The mixture was subsequently incubated with A549-ACE2 cells for 48 h after which cells were washed with PBS and lysed with Luciferase Cell Culture Lysis 5x reagent (Promega). Luciferase activity in lysates was measured using the Dual-Luciferase Assay System (Promega). The obtained relative luminescence units (RLU) were normalized to those derived from cells infected with SARS-CoV-2 pseudotyped virus in the absence of monoclonal antibodies. The half-maximal inhibitory concentrations for monoclonal antibodies (NT_50_) were determined using four-parameter nonlinear regression (least squares regression method without weighting; constraints: top, 1; bottom, 0) (GraphPad Prism).

### Biolayer interferometry (BLI) binding assays

Binding assays were performed by biolayer interferometry (BLI) using an Octet Red96e instrument (FortéBio) at room temperature as described previously.^50^ Briefly, His-tagged SARS-CoV-2 RBD proteins at 0.5 μM in 1× kinetics buffer (1× PBS, pH 7.4, 0.01% w/v BSA and 0.002% v/v Tween 20) were loaded onto anti-Penta-HIS (HIS1K) biosensors and incubated with the indicated concentrations of Fabs. The assay consisted of five steps: 1) baseline; 2) loading; 3) baseline; 4) association; and 5) dissociation. For estimating the K_D_ values, a 1:1 binding model was used.

### Expression and purification of RBD

The receptor-binding domain (RBD) of the SARS-CoV-2 spike (S) protein were cloned into a customized pFastBac vector.^51^ The RBD constructs were fused with an N-terminal gp67 signal peptide and a C34 terminal 6xHis tag. Recombinant bacmid DNA was generated using the Bac-to-Bac system (Life Technologies). Baculovirus was generated by transfecting purified bacmid DNA into Sf9 cells using FuGENE HD (Promega) and subsequently used to infect suspension cultures of High Five cells (Life Technologies) at an MOI of 5 to 10. Infected High Five cells were incubated at 28°C with shaking at 110 r.p.m. for 72 h for protein expression. The supernatant was then concentrated using a 10 kDa MW cutoff Centramate cassette (Pall Corporation). The RBD proteins were purified by Ni-NTA, followed by size exclusion chromatography, and buffer exchanged into 20 mM Tris HCl pH 7.4 and 150 mM NaCl.

### Crystal structure determination

For crystallization, purified HB148 (HB148-M4) Fab, LY-CoV1404 Fab, and SARS-CoV-2 wild-type or Omicron BA.1 RBD were mixed at an equimolar ratio and incubated overnight at 4°C. Complexes (12.5 mg/mL) were screened for crystallization with the 384 conditions of the JCSG Core Suite (QIAGEN) on our custom-designed robotic CrystalMation system (Rigaku) at Scripps Research by the vapor diffusion method in sitting drops containing 0.1 μL of protein and 0.1 μL of reservoir solution. Diffraction-quality crystals were obtained in the following conditions:

1. SARS-CoV-2 wild-type RBD/HB148/LY-CoV1404: 0.2 M di-ammonium hydrogen phosphate, 20% (w/v) polyethylene glycol 3350 at 20°C.
2. SARS-CoV-2 wild-type RBD/HB148-M4/LY-CoV1404: 0.23 M di-ammonium hydrogen phosphate, 20% (w/v) polyethylene glycol 3350 at 20°C.
3. SARS-CoV-2 Omicron BA.1 RBD/HB148-M4/LY-CoV1404: 0.2 M di-ammonium hydrogen phosphate, 12%(w/v) polyethylene glycol 3350 at 20°C.

Crystals appeared on day 3 and were harvested on day 7 by soaking in reservoir solution supplemented with 15% (v/v) ethylene glycol as cryoprotectant. The crystals were then flash-cooled and stored in liquid nitrogen until data collection. Diffraction data were collected at cryogenic temperature (100 K) at National Synchrotron Light Source II (NSLS-II) beamline 17-ID-2 and Stanford Synchrotron Radiation Lightsource (SSRL) beamline 12-1 with beam wavelengths of 0.97934 Å and 0.97946 Å, respectively. Diffraction data were processed with HKL2000.^52^ Structures were solved by molecular replacement using PHASER^53^ and one structure (PDB 7MMO) as initial model.^54,55^ Iterative model building and refinement were carried out in COOT^56^ and PHENIX,^57^ respectively.

## Data availability

The refined models have been deposited in the Protein Data Bank with accession numbers: 9ZBW, 9ZBX and 9ZBY.

## Acknowledgements

This work was supported by Calmette and Yersin scholarship from the Pasteur International Network Association (H.L.), Carl R. Woese Institute for Genomic Biology (IGB) postdoctoral fellpwship (H.L.), the Research Grants Council of the Hong Kong Special Administrative Region, China (14115125, C4002-24Y) (C.K.P.M), the Health and Medical Research Fund (22210332) (C.K.P.M), the Vallee Scholars Program (N.C.W.), the Searle Scholars Program (N.C.W.), and Howard Hughes Medical Institute Emerging Pathogens Initiative (N.C.W.).

## Author contributions

H.L., R.B., I.A.W., M.Y., and C.K.P.M.,. conceived and designed the study. H.L., Z.F., Q.W.T., C.C., A.B.G., D.C., T.J.C.T., Y.S.T., L.S., A.N., performed the experiments and data analysis. H.L., Z.F., I.A.W., M.Y., N.C.W., and C.K.P. wrote the paper and all authors reviewed and/or edited the paper.

