## Supplementary material for "Somatic evolution of a cross-reactive germline antibody that expands its breadth to neutralize new SARS-CoV-2 variants": SI figure

**Table S1. X-ray data collection and refinement statistics**

| **Data collection** | SARS-CoV-2 wild-type RBD + HB148 + LY-CoV1404 | SARS-CoV-2 wild-type RBD + HB148-M4 + LY-CoV1404 | SARS-CoV-2 Omicron BA.1 RBD + HB148-M4 + LY-CoV1404 |
| --- | --- | --- | --- |
| Beamline | NSLS-II 17-ID-2 | SSRL BL12-1 | SSRL BL12-1 |
| Wavelength (Å) | 0.97934 | 0.97946 | 0.97946 |
| Space group | P1 | C 1 2 1 | C 1 2 1 |
| Unit cell parameters |  |  |  |
| a, b, c (Å) | 85.3, 86.1, 98.8 | 195.8, 88.7, 99.9 | 196.8, 88.0, 100.8 |
| α, β, γ (°) | 115.7, 89.7, 94.4 | 90, 112.0, 90 | 90, 110.3, 90 |
| Resolution (Å) ^a^ | 50.0-3.10 (3.15-3.10) | 50.0-2.60 (2.64-2.60) | 50.0-2.73 (2.78-2.73) |
| Unique reflections ^a^ | 45,996 (2,261) | 47,927 (2,367) | 42,283 (1,794) |
| Redundancy ^a^ | 2.0 (1.9) | 4.7 (3.1) | 5.5 (3.3) |
| Completeness (%) ^a^ | 97.8 (96.7) | 98.0 (97.7) | 96.5 (82.7) |
| <I/σ_I_> ^a^ | 4.3 (1.0) | 15.4 (1.2) | 14.7 (1.3) |
| *R*_sym_^b^ (%) ^a^ | 17.6 (65.3) | 11.5 (>100) | 15.6 (>100) |
| *R*_pim_^b^ (%) ^a^ | 15.2 (58.4) | 5.7 (69.5) | 6.8 (55.9) |
| CC_1/2_^c^ (%) ^a^ | 99.6 (40.3) | 98.7 (58.8) | 98.3 (68.8) |
| **Refinement statistics** |  |  |  |
| Resolution (Å) | 48.8-3.10 | 36.1-2.60 | 40.8-2.73 |
| Reflections (work) | 39,220 | 42,405 | 34,641 |
| Reflections (test) | 1,841 | 2,109 | 2,000 |
| *R*_cryst_^d^ / *R*_free_^e^ (%) | 21.3/25.9 | 19.3/24.3 | 20.0/25.3 |
| Copies of Fab/RBD per ASU | 2 | 1 | 1 |
| No. of atoms | 15,954 | 8,159 | 8,052 |
| Fab | 12,828 | 6,420 | 6,438 |
| RBD | 3,116 | 1,558 | 1,522 |
| Ligands ^f^ | 10 | 17 | 42 |
| Water | 0 | 156 | 50 |
| Average *B-*values (Å^2^) | 69 | 50 | 50 |
| Fab | 70 | 51 | 51 |
| RBD | 65 | 49 | 48 |
| Ligands ^f^ | 75 | 51 | 54 |
| Water | - | 40 | 31 |
| Wilson *B*-value (Å^2^) | 67 | 44 | 43 |
| **RMSD from ideal geometry** |  |  |  |
| Bond length (Å) | 0.002 | 0.002 | 0.002 |
| Bond angle (^o^) | 0.50 | 0.55 | 0.52 |
| **Ramachandran statistics (%) ^g^** |  |  |  |
| Favored | 96.95 | 97.58 | 97.18 |
| Outliers | 0.05 | 0.00 | 0.00 |
| **PDB code** | 9ZBW | 9ZBX | 9ZBY |

^a^ Numbers in parentheses refer to the highest resolution shell.

^b^ *R*_sym_ = Σ*_hkl_* Σ*_i_* | I*_hkl,i_* - <I*_hkl_*> | / Σ*_hkl_* Σ*_i_* I*_hkl,i_* and R*_pim_* = Σ*_hkl_* (1/(n-1))^1/2^ Σ*_i_* | I*_hkl,i_* - <I*_hkl_*> | / Σ*_hkl_* Σ*_i_* I*_hkl,i_*, where I*_hkl,i_* is the scaled intensity of the i^th^ measurement of reflection h, k, l, <I*_hkl_*> is the average intensity for that reflection, and *n* is the redundancy.

^c^ CC_1/2_ = Pearson correlation coefficient between two random half datasets.

*^d^ R*_cryst_ = Σ*_hkl_* | *F*_o_ - *F*_c_ | / Σ*_hkl_* | *F*_o_ | x 100, where *F*_o_ and *F*_c_ are the observed and calculated structure factors, respectively.

^e^ *R*_free_ was calculated as for *R*_cryst_, but on a test set comprising ~4.7%-5.8% of the data excluded from refinement.

^f^ Bound ligands are phosphate, Tris(hydroxymethyl)aminomethane, and ethylene glycol molecules.

^g^ From MolProbity.^1^

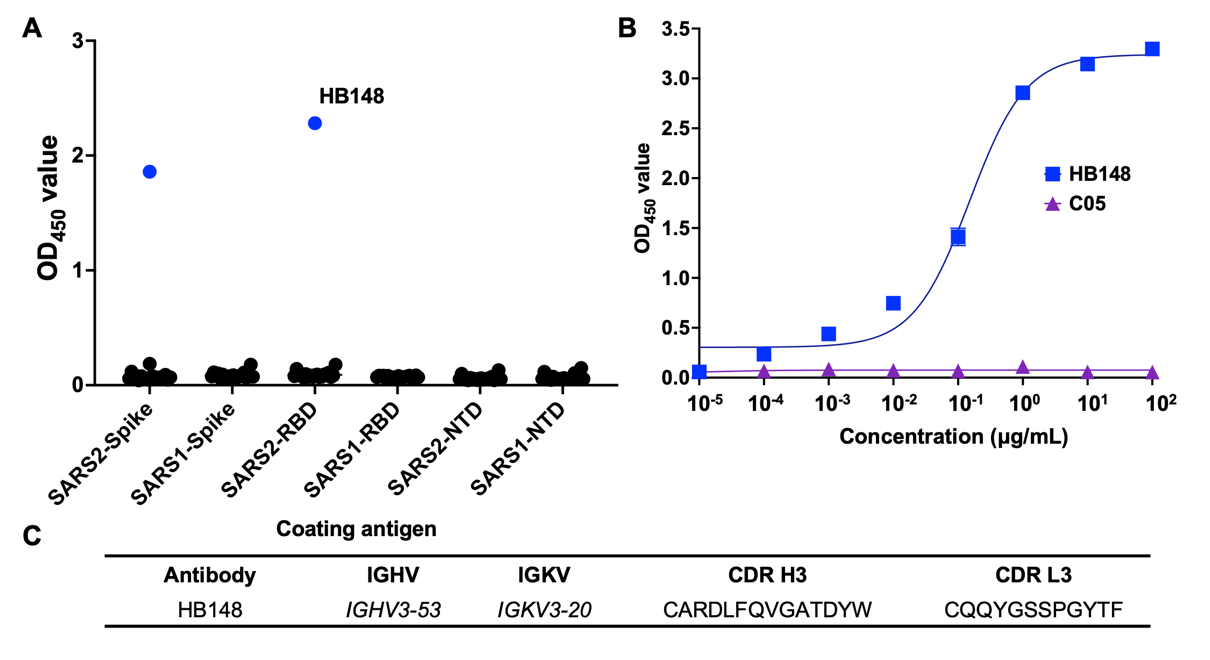

**Figure S1. Identification of human monoclonal antibodies to SARS-CoV-2 RBD. (A)** The binding activity of 24 human monoclonal antibodies against full spike, RBD, NTD protein from SARS-CoV-1 and SARS-CoV-2 was measured by ELISA. **(B)** The binding affinity of HB148 (blue) IgG against SARS-CoV-2 RBD was measured by ELISA. C05 is an influenza hemagglutinin antibody and serves as a negative control here.^2^ **(C)** Sequence information of antibody HB148 with heavy chain and light chain gene family and CDR3 amino acid sequence.

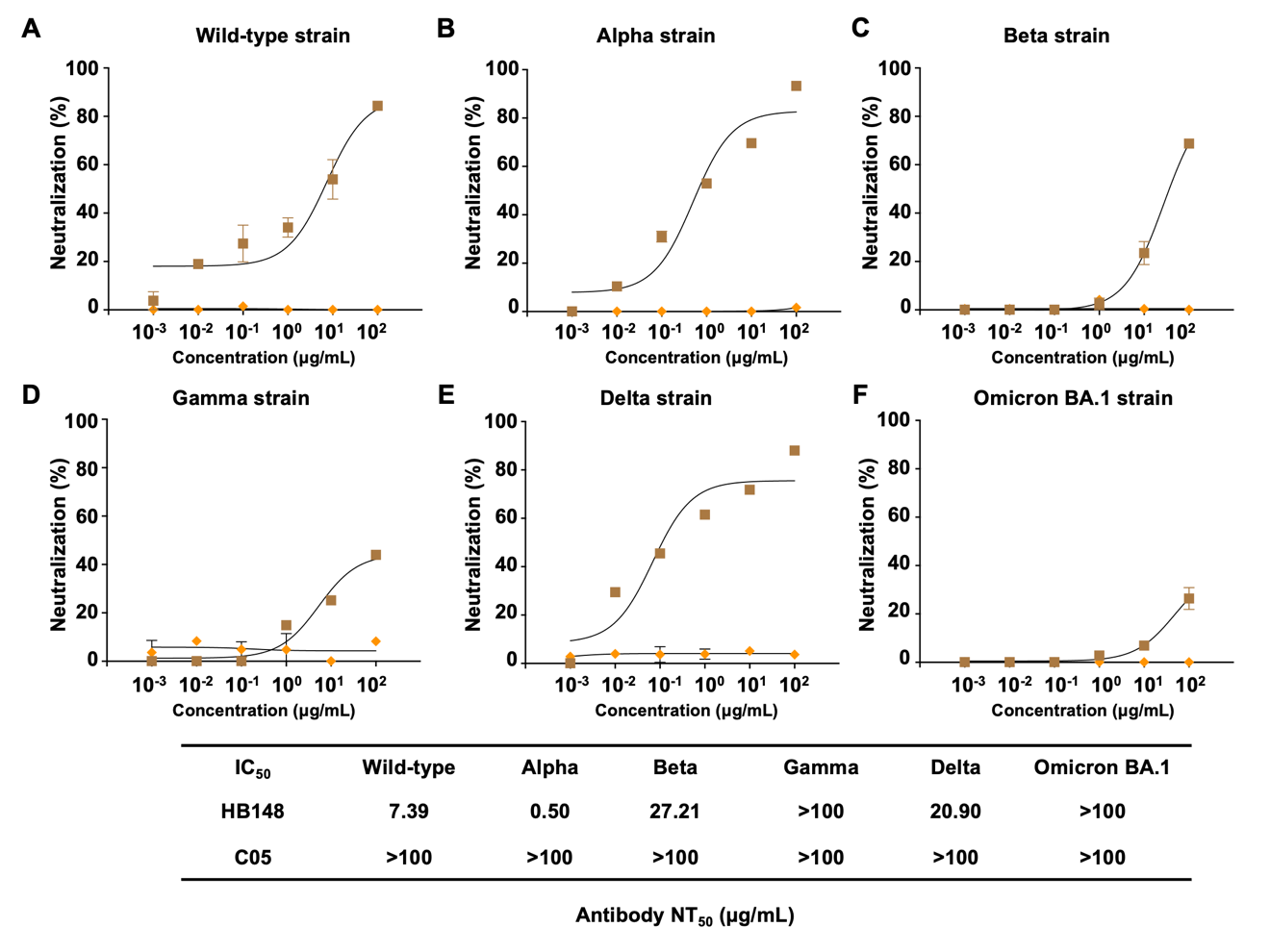

**Figure S2. Pseudovirus neutralizing ability of HB148 against different SARS-CoV-2 variants.** Neutralization activity of antibody HB148 was determined by SARS-CoV-2 pseudovirus neutralization assay against **(A)** Wild-type strain, **(B)** Alpha strain, **(C)** Beta strain, **(D)** Gamma strain, **(E)** Delta strain and **(F)** Omicron BA.1 strain. Their estimated NT_50_ values are indicated. C05 is an influenza hemagglutinin antibody and serves as a negative control here.^2^

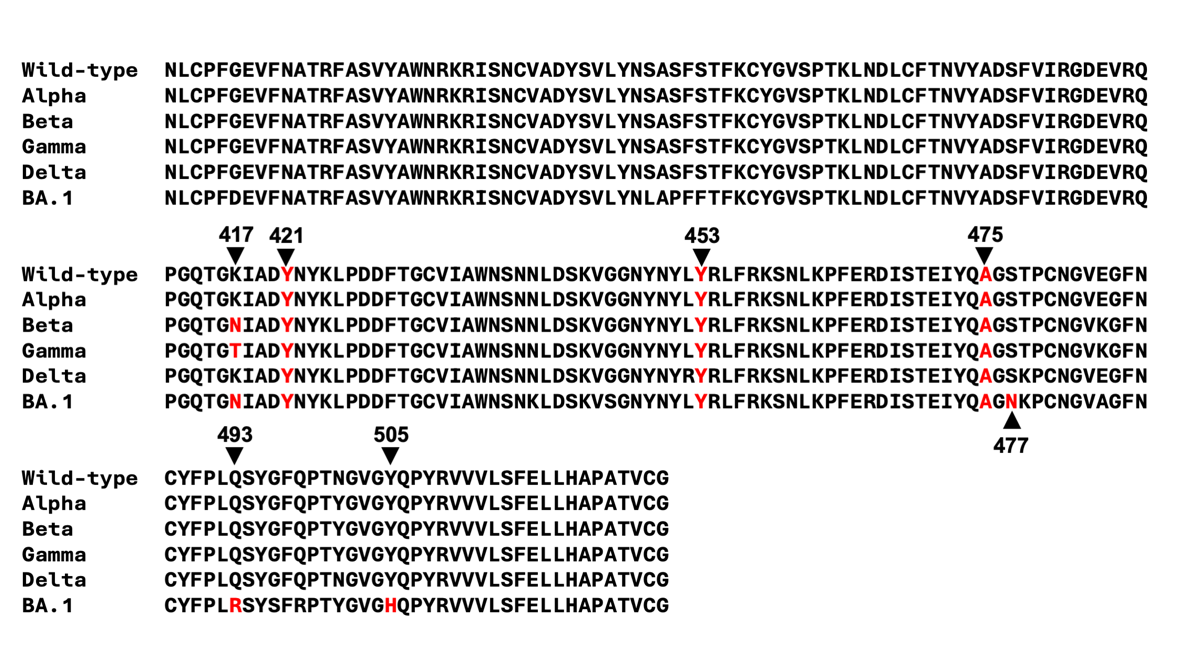

**Figure S3. Sequence alignment of RBDs from different SARS-CoV-2 variants.** Sequence alignment of RBD sequences from different SARS-CoV-2 variants was performed using MAFFT.^3^ Conserved residues among wild-type and variant strains are highlighted in red, while residues that differ from the wild-type and may contribute to escape are highlighted in blue.

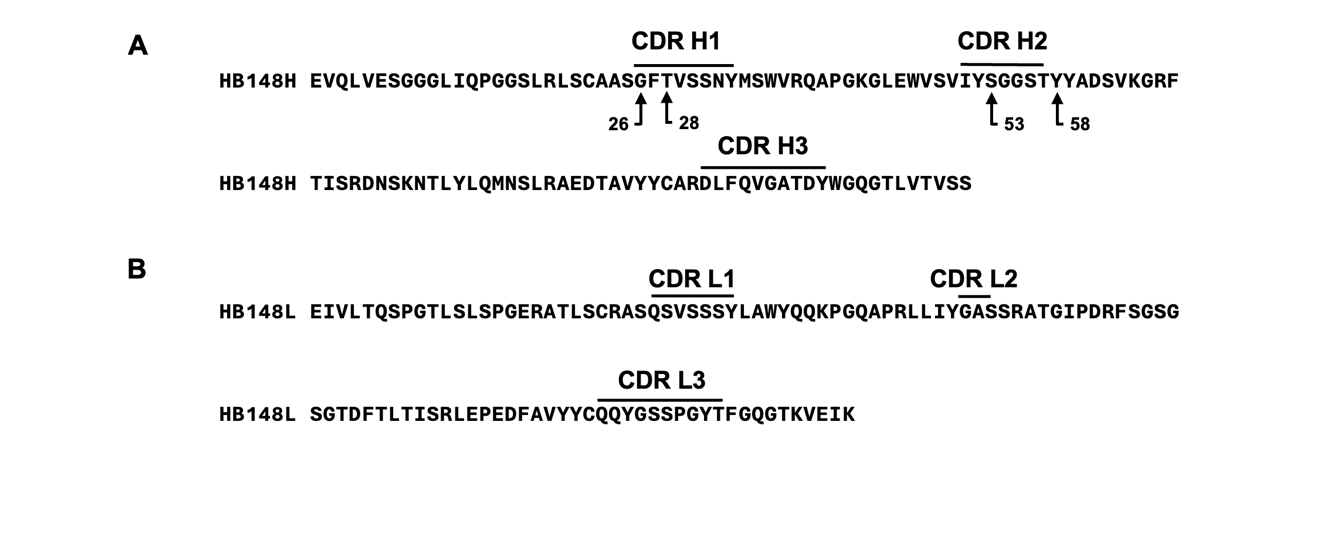

**Figure S4. Amino acid sequence information for antibody HB148 with heavy chain and light chain CDR regions highlighted.** CDR H1, CDR H2, CDR H3, and four potential somatic hypermutations in the HB148 heavy chain **(A)**, as well as CDR L1, CDR L2, and CDR L3 in the HB148 light chain **(B)**, were identified using IMGT.^4^

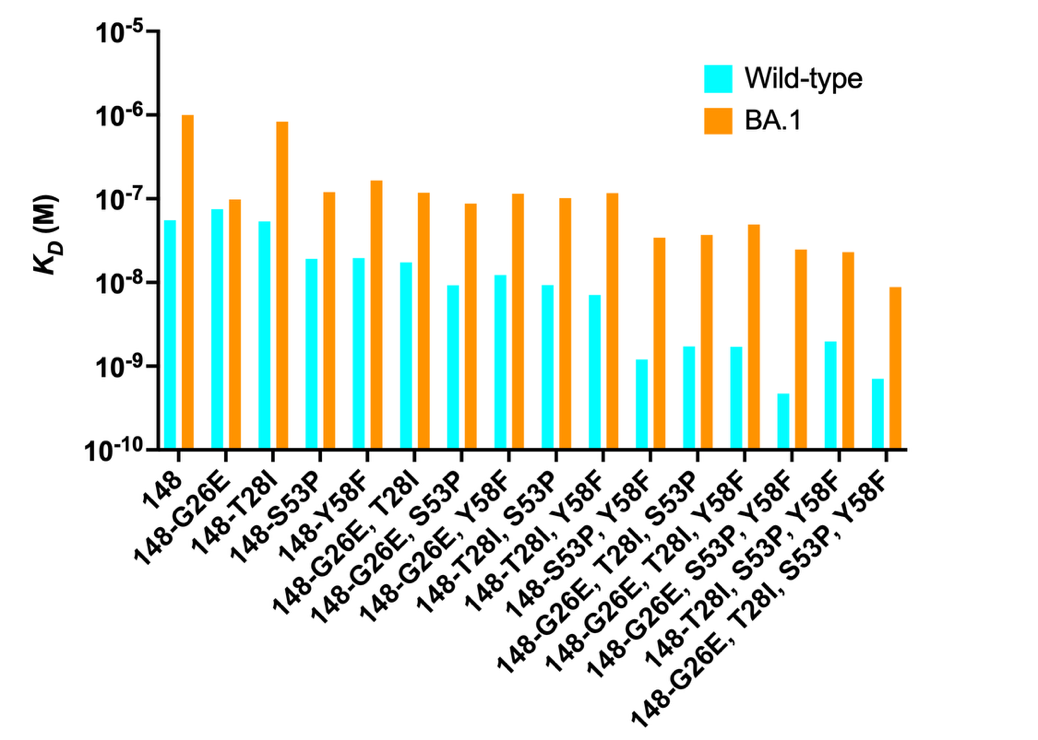

**Figure S5. Binding affinity of HB148 WT and mutants against wild-type and Omicron BA.1 RBD.** Binding kinetics of HB148 antibodies with different mutation combinations against wild-type RBD and Omicron BA.1 RBD were measured by biolayer interferometry (BLI). Y-axis represents the response.

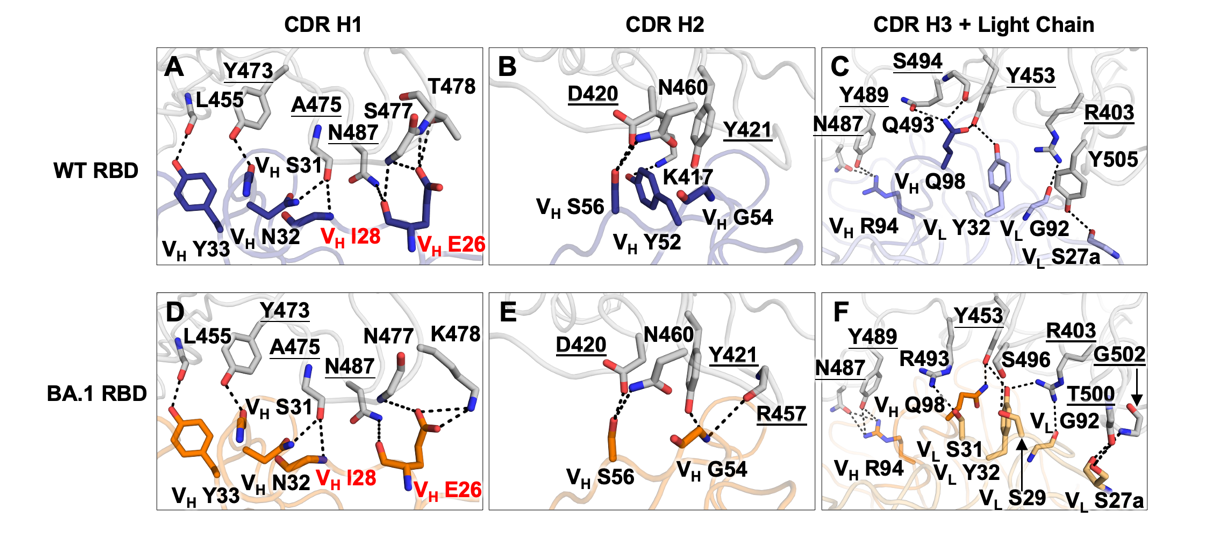

**Figure S6. Crystal structure analysis of HB148-M4 with wild-type RBD and BA.1 RBD.** Detailed molecular interactions (hydrogen bonds and salt bridges) of wild-type and BA.1 RBDs (grey backbone) with HB148-M4 are shown with V_H_ G26E and T28I labeled in red. Conserved epitope residues across VOCs are underlined.
